# Amplification of the V5 – V8 region of the 16S rRNA gene effectively speciates medically important genital tract *Lactobacillus* species in the upper female genital tract

**DOI:** 10.1101/630004

**Authors:** Jessica L. O’Callaghan, Dana Willner, Melissa Buttini, Flavia Huygens, Elise S. Pelzer

## Abstract

**Background:** The endometrial cavity is an upper genital tract site largely heralded as sterile, however, advances in culture-independent, next generation sequencing technology have revealed that this site harbours a rich microbial community which includes multiple *Lactobacillus* species. These bacteria are considered to be the most common non-pathogenic genital tract commensals. Next-generation sequencing of the female lower genital tract has revealed significant variation amongst microbial community composition with respect to *Lactobacillus* sp. in samples collected from healthy and diseased women. The aim of this study was to evaluate the ability of the 16S rRNA gene to characterize genital tract lactobacilli to species-level taxonomy.

**Methods:** Samples were interrogated for the presence of microbial DNA using two-step next generation sequencing technology to exploit the V5–V8 regions of the 16S rRNA gene and compared to standard speciation using qPCR.

**Results:** The V5-V8 region of the 16S rRNA gene has sufficient sequence variation within frequently encountered genital tract lactobacilli to allow accurate determination of relative abundance within the community, and speciation for several key community members without completing additional experimentation.

**Conclusions:** Next-generation sequencing of clinical genital tract isolates is an effective method for high throughput identification to species-level of key *Lactobacillus* sp.

**IMPORTANCE:** Human microbiome experiments, including the low biomass organs such as the upper genital tract, require the development of consensus protocols to ensure accurate comparison between such studies and our data forms an important foundation for future protocols.

This paper provides evidence to support the selection of the V5-V8 regions of the 16S rRNA gene improved *Lactobacillus* speciation using next generation sequencing technology. The choice of variable region for broad-range amplification in microbiome studies is important due to preferential primer binding associated with some genera based on nucleotide sequence patterns. By utilising the V5-V8 region, multiple species of *Lactobacillus* can be characterised with relative confidence.

## INTRODUCTION

Molecular microbiology techniques have changed our ability to identify microbial communities, revolutionizing the way we assess female genital tract microbiomes. In cultivation-dependent studies, greater than 95% of the vaginal microbiota in healthy women was classified as lactobacilli. The advent of cultivation-independent technology platforms has provided evidence to suggest that in up to two-thirds of healthy women, the lactobacilli were co-aggregated with a diverse group of microbial community members, and in some cases did not dominate (Fettweis et al., 2012; Klebanoff et al., 1991). Lactobacilli establish niche dominance through co-aggregation, competitive inhibition, production of metabolic acids, hydrogen peroxide, and antimicrobial components including bacteriocins (Amabebe and Anumba, 2018). The discovery that lactobacilli do not dominate the genital tract of all healthy women suggests that: there is redundancy in function and protection based on community membership; and all lactobacilli may not provide the same level of protection in the genital tract environment. This discovery casts doubt over the long-held view that a healthy female genital tract is characterized by a *Lactobacillus* sp.-dominant microbiota. The ability to confidently assign lower order taxonomic classification to lactobacilli is critical in advancing our understanding of the protective role played by the various species within this genus in reproductive health. The objective of this study was to examine the discriminatory power of current molecular microbiology techniques for identification of genital tract lactobacilli.

## METHODS

### Patient cohort, sample collection and genomic DNA preparation

Clinical sample cohorts were constructed as previously described (Pelzer et al., 2018). Genomic DNA was extracted from individual samples prior to pooling using a modified protocol with the Qiagen QiAMP Mini DNA extraction kit (Qiagen, Australia) as previously described (Pelzer et al., 2018).

### Details of ethical approval

All patients recruited for this study provided written informed consent. Ethical approval was obtained from the review boards of UnitingCare Health, Human Research Ethics Committee and Queensland University of Technology Human Ethics Committee.

### Next-generation sequencing

The 16S rRNA PCR assay was performed using the previously published primers, 803F (5′-TTA GAT ACC CTG GTA GTC -3′) and 1392R (5′-ACG GGC GGT GTG TRC -3′) and PCR cycling conditions (Willner et al., 2014). Fusion primers with 454 adaptor sequences were ligated to the 803F and 1392R primers to amplify the V5 and V8 regions of the 16S rRNA gene (Willner et al., 2014). PCR reactions were performed as previously described (Pelzer et al., 2018). The five frequently encountered genital tract *Lactobacillus* sp. were aligned using the SILVA database to determine the degree of variation within the V5-V8 regions of the 16S rRNA gene. The annealing site of the sequencing primers is marked on the alignment (Figure 1a).

**Figure 1a:**
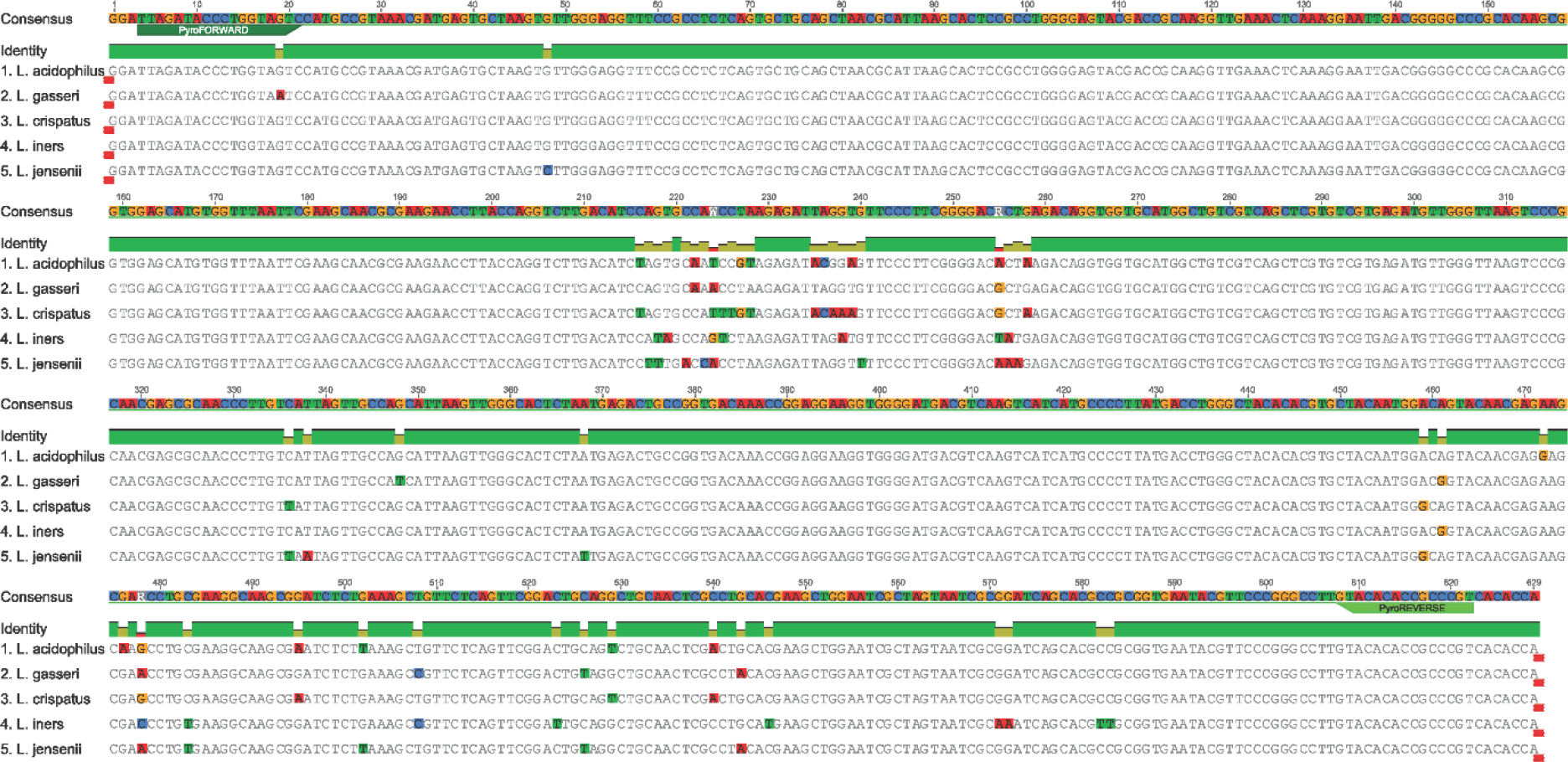
Annealing site of 454 pyrosequencing 16S rRNA primers to V5-V8 region in the *Lactobacillus* species.

### *Lactobacillus* sp.-specific quantitative real-time PCR

Quantitative real-time PCR assays were performed using previously published primer pairs (Table 1) and cycling conditions. A standard curve was generated using *L. gasseri* ATCC strain 19992. Primer annealing was confirmed using species-specific alignment of the five *Lactobacillus* sp. interrogated in this study (Figure 1b).

**Table 1.** Sample abbreviations

| Sample Abbreviations | Full Sample Name |
| --- | --- |
| DGC | dysmenorrhea progestin effect endocervix |
| DSC | dysmenorrhea secretory endometrium |
| MGC | menorrhagia progestin effect endocervix |
| DPC | dysmenorrhea proliferative endocervix |
| DGE | dysmenorrhea progestin effect endometrium; |
| DPE | dysmenorrhea proliferative endometrium |
| MGE | menorrhagia progestin effect endometrium |
| MSE | menorrhagia secretory endometrium |
| MSC | menorrhagia secretory endocervix |
| MPE | menorrhagia proliferative endometrium |
| MPC | menorrhagia proliferative endocervix |
| VIC | virgo intacta |

**Figure 1b:**
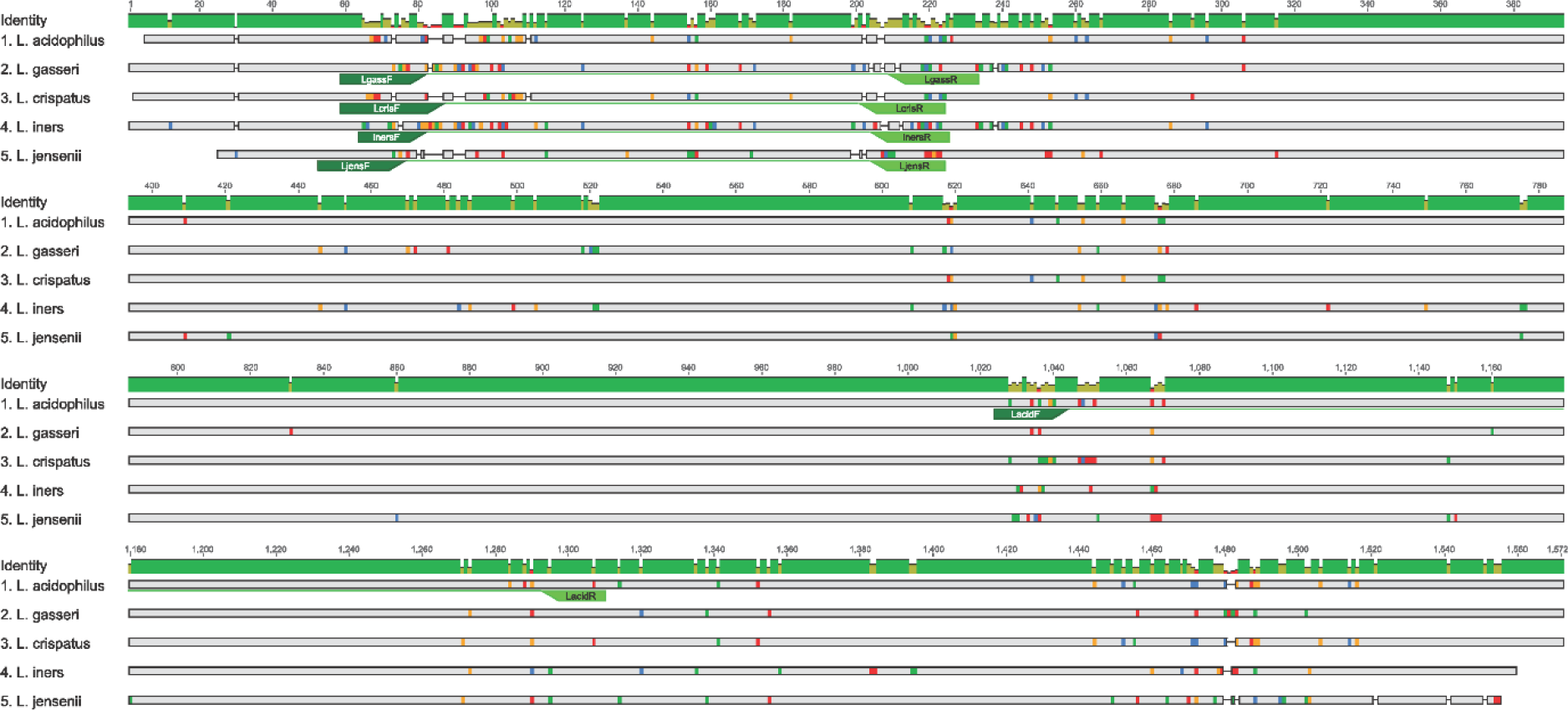
Annealing site of species specific primers (Table 2) in the *Lactobacillus* species which underwent qPCR.

**Table 2.**
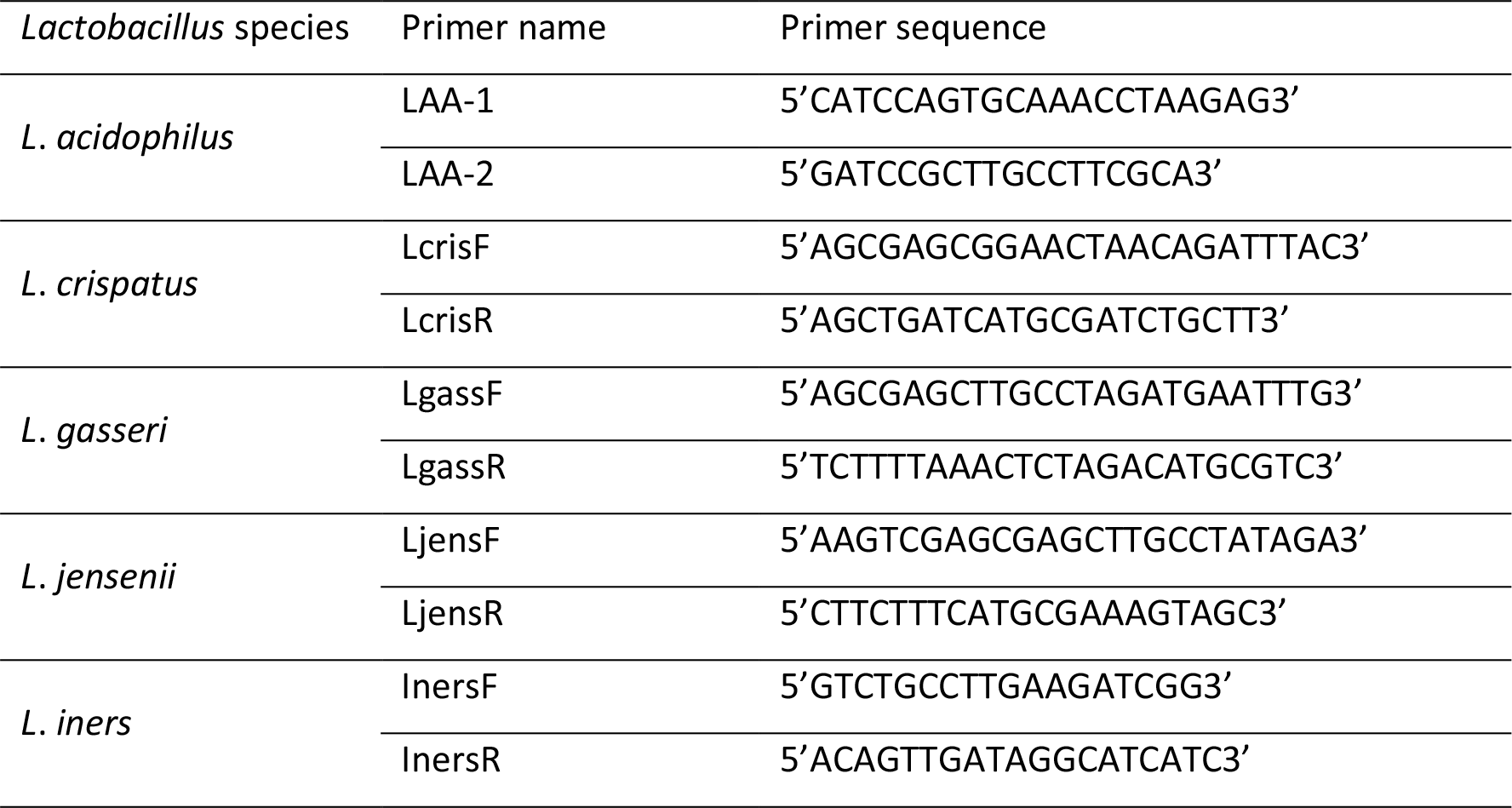
16S rRNA *Lactobacillus*-species specific primers (Ma et al., 2013)

### Taxonomic classification

Sequence clustering and operational taxonomic unit (OTU) selection was performed using a modified version of CD-HIT-OTU-454 which does not remove singleton clusters (Liu et al., 2011). Taxonomy was assigned to representative sequences by comparison to the latest build of the Greengenes database using BLAST, and OTU tables were constructed from the output using a custom Perl script (McDonald et al., 2012).

### *Lactobacillus* Phylogenetic Trees

Full-length 16S rRNA sequences for *Lactobacillus spp.* (accession numbers: AB680529.1, AB690249.1, AB668940.1.1, AB008203.1.1, AB425941.1.1, AB008206.1, AF243169.1, AF243167.1, CP018809.253324, CP018809.1516019, CP018809.1347636, CP018809.500868, AB547127.1, AB517146.1, AB932527.1, AB008209.1, HZ485829.7, LG085736.7 LF134126.7, LG104504.7), *Pediococcus pentosaceus* (accession numbers: AB018215.1 and AB362987.1), and *Bacillus subtilis* (accession numbers: AP012496.9810 and AP012496.30276) were downloaded from the SILVA database using the web interface (www.arb-silva.de). Sequences were aligned using ClustalW (Thompson et al., 1994) with the default settings. MEGA7 (Kumar et al., 2016) was used to generate the best-known maximum likelihood (ML) tree using a Jukes-Cantor model and 1000 bootstrapping iterations. The ML tree was visualised and edited within FigTree (http://tree.bio.ed.ac.uk) and Adobe Illustrator (Figure 4).

To generate the V5-V8 region phylogenetic tree the same full length 16S rRNA sequences from above were imported into Geneious along with two *Escherichia coli* sequences downloaded from SILVA (accession number: AB045730.1 and AB045731.1). Sequences were aligned using standard Geneious alignment and trimmed to include the variable regions V5-V8 (nucleotides 751 – 1300 for the *E.coli* sequence). Trimmed sequences were then aligned using ClustalW with default settings and imported into MEGA7 and a tree was constructed and edited same as above (Figure 4).

### Hierarchical clustering

A dissimilarity matrix was generated based on the relative abundances of *Lactobacillus spp.* in the pyrosequenced and qPCR analysed samples using the vegdist function in the vegan package in R with the Bray-Curtis dissimilarity metric (J. et al., 2018). Hierarchical clustering was performed using the hclust function in R with ‘average’ linkage (UPGMA) (Team, 2013). Clustering and relative abundances were visualized in a heatmap with associated dendrogram using the heatmap.2 function from the R package ggplots (Warnes et al., 2005).

## RESULTS

### NGS resolution of genital tract *Lactobacillus* sp. OTUs to genus and species

Ten OTUs were attributed to *Lactobacillus* sp. (*Lactobacillus* sp. genus level (n = 2), *L. crispatus* (n = 2), *L. iners* (n = 3), *L. intestinalis* (n = 1), *L. jensenii* (n = 1) and *L. vaginalis* (n = 1)). The majority of *Lactobacillus* sp. OTUs (8/10) were resolved to the genus and species level exploiting the V5-V8 regions of the 16S rRNA gene.

### *Lactobacillus* species-specific quantitative real-time PCR assay comparison to NGS output

The species-specific quantitative real-time PCR assays confirmed the identity and relative abundance of *L. crispatus*, *L. jensenii* and *L. iners* in clinical genital tract samples. Two of the species that underwent pyrosequencing identification were not included due to low abundance (*L. intestinalis* and *L. vaginalis*).

The abundance of the five commonly encountered species (*L. crispatus*, *L. gasseri*, *L. iners*, *L. jensenii* and *L. acidophilus*) were then compared between the qPCR and the 454 pyrosequencing (Figure 2). *L. crispatus* dominated the *Lactobacillus* community in most samples. All samples displayed similar abundance profiles with enriched lower abundance species including *L. jensenii*, *L. iners* and *L. acidophilus* exposed by qPCR (Table 3, Figure 2). The V5-V8 region of the 16S rRNA gene was not as effective in distinguishing *L. acidophilus* from *L. gasseri* using the primers published by Ma *et al.* (2013) due to high sequence homology between these two species in the V5-V8 variable region.

**Figure 2:**
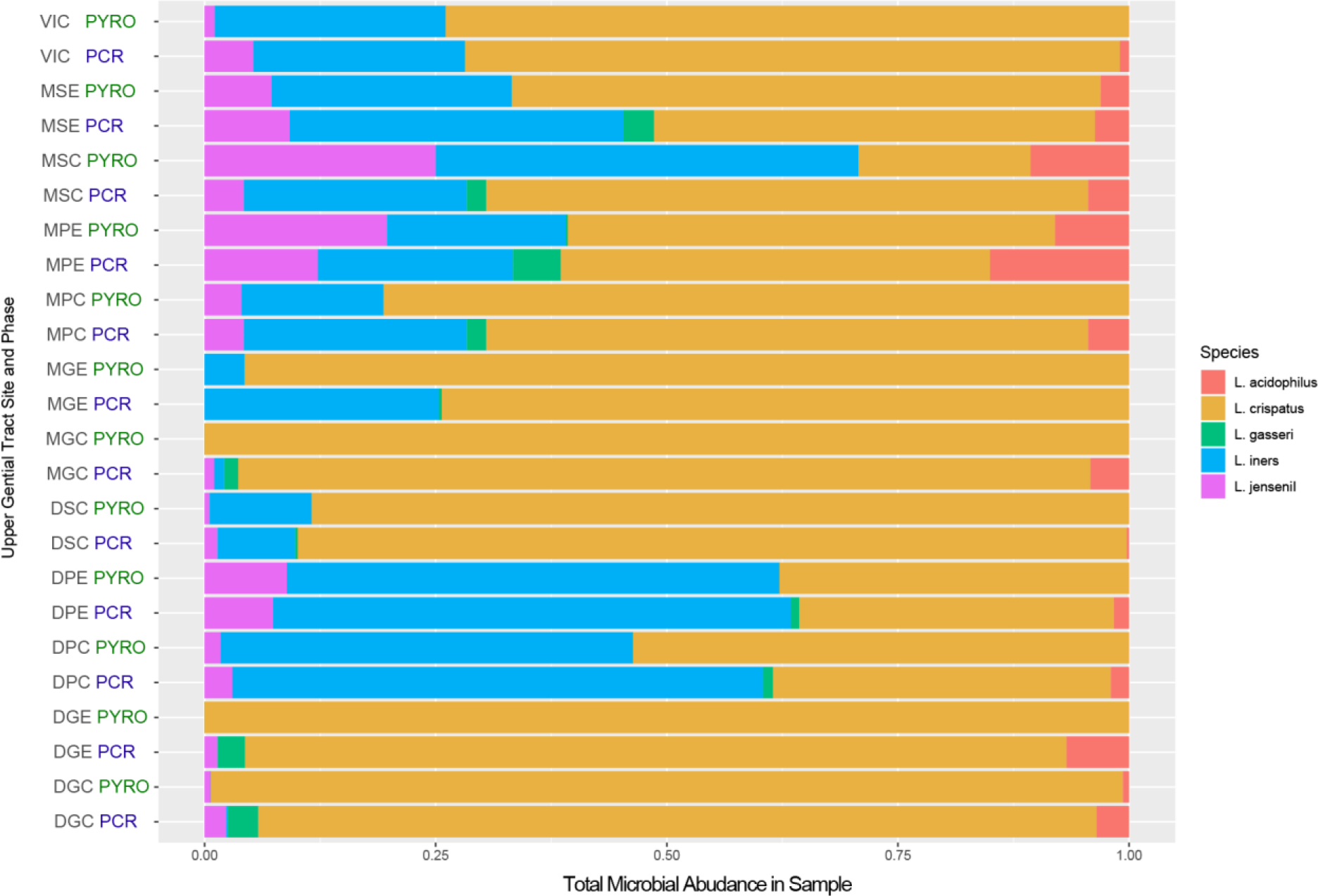
Comparison of relative microbial abundance all sites and phases between 454 pyrosequencing of the V5 - V8 region and species specific Lactobacillus qPCR.

**Figure 3:**
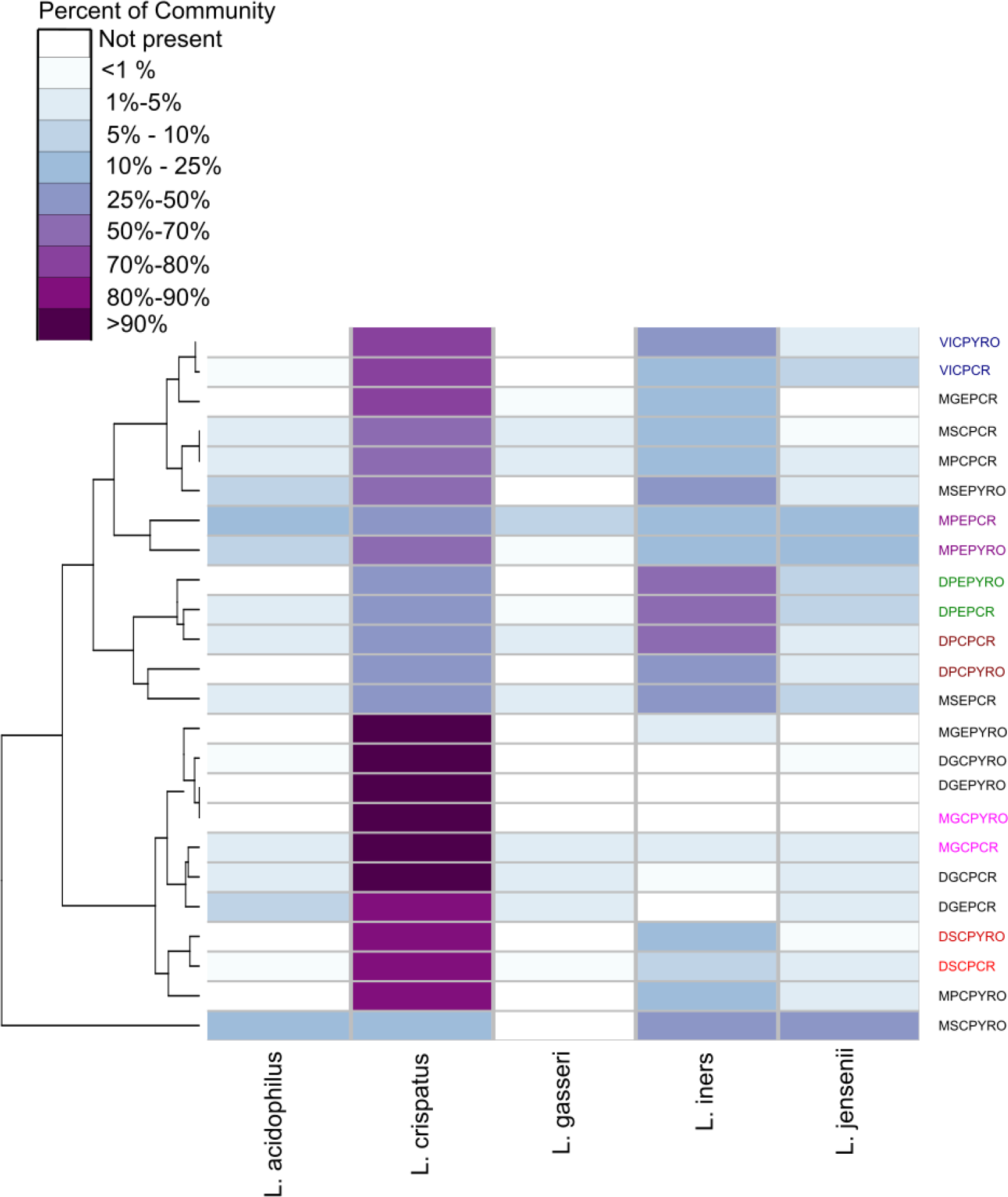
Hierarchical clustering/distance ordination quantifying similarities between qPCR and PYRO pairs. *Lactobacillus* relative abundance distribution and hierarchical clustering in pyrosequenced and qPCR analysed samples. The heatmap shows relative abundances of detected *Lactobacillus* species in each sample, with columns organised by relative positions in the dendrogram. Some subsets of paired pyrosequenced and qPCR samples have been coloured to highlight clustering.

**Figure 4a:**
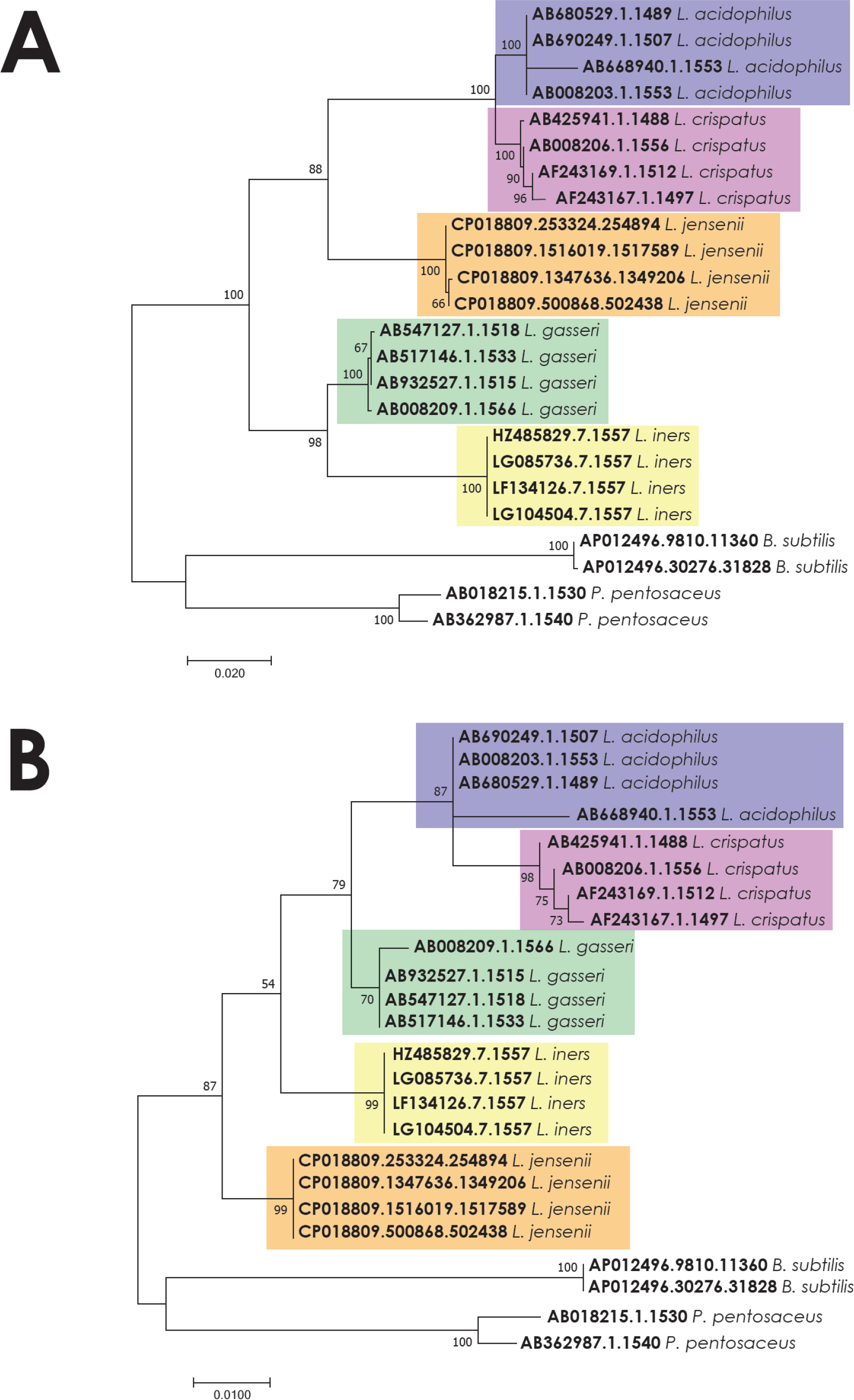
Full length and region specific 16S rRNA gene phylogeny for key genital tract lactobacilli. ML tree derived from full-length and trimmed (V5-V8 region) 16S rRNA sequences of the multiple species of *Lactobacillus* identified by pyrosequencing and 16S qPCR. Accession numbers for reference sequences are provided in the methods. *Pediococcus pentasaecus* and *Bacillus subtilis* were included as outgroups. Branch support values are based on 1000 bootstrap repetitions.

**Table 3:** Unique to species SNPs from the 16S rRNA gene variable regions.

| Species | 16S rRNA gene variable Region |  |  |  |  |  |  |  |  |
| --- | --- | --- | --- | --- | --- | --- | --- | --- | --- |
|  | V1 | V2 | V3 | V4 | V5 | V6 | V7 | V8 | V9 |
| *L. iners* | 5 | 17 | 0 | 2 | 0 | 0 | 1 | 2 | 4 |
| *L. gasseri* | 4 | 0 | 2 | 0 | 0 | 0 | 1 | 0 | 1 |
| *L. acidophilus* | 2 | 4 | 0 | 3 | 2 | 0 | 0 | 2 | 0 |
| *L. crispatus* | 2 | 1 | 0 | 1 | 3 | 2 | 0 | 0 | 0 |
| *L. jensenii* | 6 | 6 | 0 | 1 | 1 | 5 | 1 | 0 | 5 |

### *Lactobacillus* phylogeny

The phylogenetic tree constructed using the full-length16S rRNA gene sequences of the *Lactobacillus* sp. described in this study indicated that *L. acidophilus* and L. *crispatus* appear more closely related than *L. acidophilus* and *L. gasseri*. When comparing only the V5-V8 region, however, *L. gasseri* clusters closer to *L. acidophilus* (Figure 4). The heat map confirmed that *L. crispatus* and *L. iners* dominated the microbial communities in samples analyzed in this study and samples were more likely to cluster based on *Lactobacillus* community dominance, than the patient history (dysmenorrhea or menorrhagia), the anatomical site of collection (endometrium or cervix), or the analysis technique (qPCR or pyrosequencing).

## DISCUSSION

Sequencing of the V5-V8 region of the 16S rRNA gene improves the discriminatory power for speciation of dominant genital tract lactobacilli. This study examined the different bacterial communities within the upper genital tract of women, reporting changes in the bacterial community composition of lactobacilli. Consistent with previous studies, *L. crispatus* and *L. iners* were the most abundant lactobacilli in the samples tested in this study.

Sequencing technologies frequently often only report the presence of lactobacilli at genus-level. Studies exploiting some regions of the 16S rRNA gene fail to discriminate lactobacilli beyond higher order taxonomic classification due to limited sequence variation. Therefore, some studies have reported that lactobacilli as a genera are: positively correlated with healthy pregnancy outcomes including successful implantation and delivery at term; and form abundant community members in cases of adverse pregnancy outcomes including recurrent implantation failure and preterm birth (Franasiak et al., 2016; Moreno et al., 2016; Onderdonk et al., 2008; Tao et al., 2017). Our research design enabled us to overcome the shortfalls commonly associated with genus-level identification. Similar results can be observed in molecular studies characterising the female genital tract when multiple variable regions of the 16S rRNA gene were sequenced (Fettweis et al., 2012; Graspeuntner et al., 2018; Madhivanan et al., 2014; Miles et al., 2017). Van Der Pol *et al.* (Van Der Pol et al., 2019) reported that the choice of 16S rRNA reference sequence database and sample sequence clustering parameters are equally as important as the choice of variable region for amplification characterising microbial community members to lower orders.

There is no doubt that sequencing the conserved 16S rRNA gene has improved our understanding of extant biodiversity in human microbial communities and is critical for understanding the impact of low-abundance community members on health and disease. However, there is no consensus best practice for microbiome studies, and significant variability exists between sample collection and storage methods, DNA extraction, universal primer selection, and sequencing platform and data analysis software (Pollock et al., 2018). Characterization of microbial communities using the 16S rRNA gene have been hampered by inherent differences generated in community profiles when sequencing different hypervariable regions, short read lengths, and taxonomic classification difficulties due to limited resolution for closely related species (Poretsky et al., 2014). Sequencing technologies have been used to interrogate the genital tract microbial community in reproductive-aged women but most fail to resolve the isolates to species-level. Consequently, more recent efforts have focused on sequencing multiple variable regions of the gene with amalgamation of all data into a single profile (Fuks et al., 2018). Very current research focuses on removing bias associated with sequencing component variable regions by using full-length gene sequencing (Callahan et al., 2016). The need to characterise the full-length 16S rRNA gene is further required as exhibited by the change in *L. gasseri* clustering when comparing the full-length gene to the V5-V8 region. Within this study, these species were not able to be distinguished from each other using this region alone.

The significance of our research is highlighted by studies confirming that *L. iners* does not protect against preterm birth and is frequently reported as an abundant community member in women with bacterial vaginosis (Madhivanan et al., 2014; Petricevic et al., 2014). Further, significant differences between lactobacilli in term compared to preterm deliveries have not been reported for all studies (Amabebe and Anumba, 2018; Romero et al., 2014). Within our study we are able to identify the bacteria to a species level using pyrosequencing reads with relative confidence. One limitation of this study is the relatively small sample size.

Collectively our research confirms what other studies have shown, that health and disease may depend on species and strain-level differences for prominent community members at a given anatomical niche (Kraal et al., 2014). It is clear that additional discriminatory power is required to resolve lower order classifications using current sequencing methods. This current study confirms that speciation of key genital tract *Lactobacillus* sp., capable of modulating reproductive health is possible when the appropriate region of the 16S rRNA gene is interrogated.

## CONCLUSION

Studies characterizing microbial communities in the female genital tract report inconsistent results when assessing dysbiosis as a cause of reproductive pathology. Our work provides evidence for the impact of primer selection on evaluating the biological significance of shifts in community taxa. Careful experimental design should include a comparative analysis of microbial community profiling data generated by interrogation of multiple variable regions to the 16S rRNA gene to ensure that species abundance and diversity are accurately reflected.

## List of abbreviations

ATCC: American type culture collection
DGC: dysmenorrhea progestin effect endocervix
DGE: dysmenorrhea progestin effect endometrium
DNA: deoxyribonucleic acid
DPC: dysmenorrhea proliferative endocervix
DPE: dysmenorrhea proliferative endometrium
DSC: dysmenorrhea secretory endocervix
DSE: dysmenorrhea secretory endometrium
HRM: high resolution melt
MGC: menorrhagia progestin effect endocervix
MGE: menorrhagia progestin effect endometrium
MPC: menorrhagia proliferative endocervix
MPE: menorrhagia proliferative endometrium
MSC: menorrhagia secretory endocervix
MSE: menorrhagia secretory endometriumx
OTU: operational taxonomic unit
rRNA: ribosomal ribonucleic acid
VIC: virgo intacta

## ACKNOWLEDGEMENTS

The authors wish to thank Wesley Hospital theatre staff who facilitated collection of the genital tract samples. We wish to acknowledge the Australian Centre for Ecogenomics for 454 pyrosequencing and Professor Philip Hugenholtz and Dr Fiona May. This work was performed in the John and Wendy Thorsen Women’s Health Laboratory.

## CONTRIBUTION TO AUTHORSHIP

JOC: designed and completed bioinformatics analyses, contributed to the analysis and interpretation of the data, and contributed to the writing of the manuscript.

DW: designed and completed bioinformatics analyses, contributed to the analysis and interpretation of the data, and contributed to the writing of the manuscript.

MB: conceived and designed the project, performed collection of clinical specimens and contributed to the writing of the manuscript.

FH: designed and completed the qPCR experiments, contributed to the analysis and interpretation of the qPCR data and contributed to the writing of the manuscript.

EP: conceived and designed the project, completed tissue processing, DNA extraction and 16S PCR experiments, contributed to the analysis and interpretation of the data, and drafted significant parts of the work.

## CONFLICT OF INTEREST STATEMENT

The authors declare that there are no conflicts of interest.

## FUNDING STATEMENT

Funding for this project was awarded by the Wesley Research Institute (Grant number 2011-12). The funding body played no role in conducting the research or preparing the manuscript.

